# Genome-wide association study in Heterogeneous Stock rats identifies genetic loci associated with aversion-based learning and cocaine aversion

**DOI:** 10.64898/2026.08.07.743619

**Authors:** Zachary Tatom, Maya Eid, Thiago Missfeldt-Sanches, Apurva S. Chitre, Gavrila Ang, Kendra S. Ziegler, Beverly Peng, Elaine Keung, Khai-Minh Nguyen, Katarina Cohen, Yizhi Wang, Riyan Cheng, Denghui Chen, Benjamin B. Johnson, Oksana Polesskaya, Tom C. Jhou, Abraham A. Palmer

## Abstract

Addiction is a complex and heritable trait which progresses through several developmental stages, each of which is presumably influenced by multiple partially overlapping genetic factors. Cocaine initially produces rewarding effects, followed by aversive effects including anxiety, craving, anhedonia, and withdrawal. These aversive effects have been suggested to contribute to the etiology of cocaine use disorders (CUD), as repeated exposure is thought to desensitize the rewarding effects and sensitize the aversive effects through a process involving both aberrant reward-based learning and aberrant avoidance-based learning. We examined the genetic basis of aversion learning using both food-based and cocaine-based behavioral assays in outbred Heterogenous Stock (HS) rats. A total of 1,074 HS rats (35.3% male) underwent runway operant cocaine-seeking, food-based progressive ratio and punishment testing, and locomotion testing. These phenotypes were significantly heritable (with *h*^2^ estimates as high as 0.307) and identified significant (*p* < 0.05) genetic loci related to avoidance-based learning including from the punishment task on Chromosomes 2, 3, 5, and 6, and the cocaine-operant runway latency task on Chromosome X. 172 positional candidate genes were identified from significant and suggestive loci, including *Cdh10*, *Cdh12*, *Cdh18* which have previously been associated with smoking initiation from human GWAS, *Adcy3*, *Cfap206*, and *Drc1* which are associated with primary neuronal cilia, as well as SNPs associated with novelty-related and social interaction phenotypes in independent samples of HS rats. Our results suggest that these aversion learning phenotypes are themselves complex heritable traits influenced by multiple genetic loci, which may pleiotropically affect other aspects of addiction biology.

## Introduction

Cocaine use presents a significant threat to public health, with notable increases in cocaine production, sale, and use observed in recent years. The United Nations estimates that 25 million individuals used cocaine in 2023, compared to 17 million in 2013^1^. Similarly, deaths attributable to cocaine overdose have increased from 3,822 in 1999 to 29,449 in 2023^2^, with additional deaths attributed to polysubstance overdoses that include cocaine^3^. Cocaine use, abuse, and dependence are highly heritable with twin-based heritability estimates ranging from 0.39 to 0.79 depending on the trait and population sample^4–6^. Despite cocaine use disorder (CUD)’s heritability, the genetic mechanisms underlying CUD are not well-understood, in part because of the genetic complexity and pleiotropy but also due to the complex etiology of addiction. Development of CUD progresses through multiple stages at which individual variability can contribute to vulnerability, such as initial exposure, escalation of use, or withdrawal.

Much of the research into the development of a CUD focuses on aberrant reward learning, with the dopaminergic reward system thought to be involved in the initial rewarding effects. However, cocaine produces an initial rewarding or euphoric phase followed by a period of aversive effects. This pattern is the basis of the Opponent-Process Theory of drug administration^7^. Dysregulation of these aversive effects has also been suggested to contribute to the progression of CUD, as repeated use is thought to desensitize the rewarding effects and sensitize the aversive effects of cocaine^8–10^. Cocaine is amenable to studying these aversive effects, as they occur quickly and reliably in rodent models about 15 minutes after exposure.^8,11^

To better understand the genetic architecture of CUD, human genome-wide association studies (GWAS) have been employed to identify variants and genes associated with dependence^12^ and CUD^13,14^. These human GWAS have identified a limited number of genes which have not yet replicated across different samples or led to successful therapeutic targets, possibly due to small sample sizes (particularly small numbers of cocaine-exposed individuals) and significant environmental effects. Animal model GWAS of cocaine are a valuable complement to human GWAS of CUD because they allow for control over these environmental factors in an experimental setting.

As an outbred model, Heterogeneous Stock (HS) Rats, have high levels of genetic diversity and recombination events which allow for fine-mapping genetic studies of complex traits^15,16^, including CUD-related traits such as cocaine self-administration^17–19^. These studies have identified novel loci associated with cocaine addiction-like behaviors while also replicating one gene, *Trak2*, previously associated with CUD in human GWAS^18,20^.

In this study, we hypothesized that GWAS of cocaine aversion and other food-based aversion-related behaviors in HS rats enables identification of new loci associated with avoidance-based learning, a key factor in the etiology of SUD. We assayed 1,074 HS rats for cocaine aversion using a runway operant cocaine-seeking task followed by food-based testing for punishment resistance and progressive ratio. We then genotyped these animals and conducted GWAS to identify associated loci.

## Methods

### Animals

Heterogeneous Stock Rats [NMcwi:HS #2314009 (RRID:RGD_2314009)] were obtained from a breeding colony at the Medical College of Wisconsin. All rats weighed 250–350g upon delivery from vendor and were individually housed in standard ventilated cages in a vivarium maintained at 22° C, and with a 12/12 h light/dark cycle (lights on at 6:00am), with food and water provided *ad libitum*, unless otherwise stated. Procedures conformed to the National Institutes of Health *Guide for the Care and Use of Laboratory Animals*, and institutional protocols approved by the MUSC Institutional Animal Care and Use Committee (IACUC). All animals were tested during the light phase.

### Surgery

Rats were anesthetized with isoflurane and fitted with indwelling intravenous catheters inserted into the right jugular vein and subcutaneously passed to a guide cannula exiting the animal’s back. After surgery, catheter patency was maintained via daily flushing with 0.05 ml of Taurolidine Citrate Solution. Animals recovered for 7 days prior to behavioral testing. Catheter patency was assessed periodically through observation of the loss of the righting reflex after i.v. injection of methohexital (Brevital, 2.0 mg/kg/0.1 ml). Rats that were unresponsive to Brevital were implanted with a new catheter using the left jugular vein.

### Behavioral testing

#### Runway Operant Cocaine-Seeking

The details of these methods have been described in a previous work [9]. Briefly, the runway consists of two opaque plastic compartments (“start” and “goal”, 25x10x17 cm length/width/height) connected by a 170cm long corridor with motorized doors between the start/goal boxes and corridor. On test days, rats were tethered to an intravenous line, then placed into the start box for 30 seconds, after which doors were opened to allow free exploration of the apparatus. Entry into the goal box caused doors to close and a syringe pump to deliver cocaine (0.75mg/kg iv over 10 seconds, same dose as in CPP and electrophysiological recording). Rats then remained in the goal compartment for 5 minutes before being returned to their home cage. Failure to enter the goal compartment after 15 minutes resulted in a timeout, in which case animals were gently nudged by the experimenter into the goal box and retained there for 5 minutes, ensuring that all animals received the same exposure to both cocaine and the goal compartment. Prior to cocaine trials, animals received habituation trials in which animals freely explored the entire apparatus without cocaine. Rats received a minimum of two trials, and continuing until two consecutive habituation trials produced latencies below 60 seconds. Animals then received 7 cocaine trials, at most twice daily and at least 4 hours apart, during the light portion of the 24-hour diurnal lighting cycle, over a period of 4-5 days. In all trials, we tabulated both the latency to reach the goal box (after exiting the start box), as well as the number of run reversals, defined as a change in run direction spanning a minimum of two photobeams (∼12 inches).

#### Progressive Ratio and Punishment Resistance

Training procedures are identical to those we previously described [13]. Unlike the runway operant cocaine-seeking task, these assays rely on food as a reward. Animals are food restricted to 85% of initial body weight, trained to lever press for food pellets (45mg) until stable response rates were reached, then tested either on the PR task with progressively increasing effort requirements (lever presses required per pellet) [16] or progressively increasing punishment (i.e. a foot shock of increasing intensity presented 1 second after food pellet delivery, whose magnitude increases about 25% every 3 trials). In the PR task, rats time out if two minutes passed without a lever press. In the punishment task, a timeout is defined by 3 consecutive trials with >30 seconds passing without a lever press. For both tasks, the breakpoint is the last successfully completed ratio or shock intensity endured before timeout.

#### Locomotion

Exploratory locomotion in a novel environment was recorded in operant chambers (12” wide, 9.5” deep, and 11.5” high) placed inside sound-attenuating cabinets (Med-Associates). Movement within the chamber was detected via interruptions of an array of 4 photodetector/emitter pairs. Locomotor counts were averaged over two 30-minute sessions on consecutive days.

### Genotyping

Spleen samples were collected postmortem for genotyping 10.17504/protocols.io.6qpvr665ovmk/v1), and DNA was isolated using the Beckman Coulter DNAdvance Kit at the University of California San Diego 10.17504/protocols.io.8epv59reng1b/v1). All samples were normalized and processed in a randomized order prior to library preparation 10.17504/protocols.io.261genw5dg47/v1), and multiplexed sequencing libraries were prepared using the iGenomX RipTide kit 10.17504/protocols.io.j8nlkkm85l5r/v1). Final QC was performed on sequencing libraries; sequencing was performed using an Illumina NovaSeq 6000 10.17504/protocols.io.yxmvmnw29g3p/v1). Reads were demultiplexed using fgbio v1.3.0 (http://fulcrumgenomics.github.io/fgbio/) before trimming adapters using BBDuk v38.94 (https://sourceforge.net/projects/bbmap/) and quality trimming using Cutadapt v4.1.68. Reads were aligned to the rat reference genome mRatBN7.2 from the Rat Genome Sequencing Consortium (GCA_015227675.2 GCF_015227675.2) using BWA-mem v0.7.17.

Sequences were then used to construct haplotypes and impute biallelic SNP genotypes using STITCH v1.6.6.70 for a total of 7,347,327 SNPs. We then removed any SNPs with low imputation quality scores (INFO <0.9; 2,609,890 SNPs removed). We additionally removed all SNPs with high missing rates (missing rate > 0.1; 22,686 removed), low minor allele frequencies (MAF <0.005; 1,905,988 removed), and extreme deviations from Hardy–Weinberg Equilibrium (HWE p < 1e-10; 2,941 removed). Some SNPs were removed based on more than one of these criteria. The remaining 5,416,497 SNP genotypes were used for all further analyses. A total of 1,174 HS Rats were genotyped with 1,074 animals passing sample-level QC metrics (reported sex and chromosomal sex mismatches, high missingness levels in genotyping, or missing data).

### Phenotype Processing, Heritability Estimation, and Genetic Correlation

Phenotypes used for analysis included body mass and data from two locomotor sessions. For the food-seeking task, Progressive Ratio was calculated as highest number of consecutive presses emitted for a single 45mg food pellet. From the punishment resistance task, Punishment was measured as the highest footshock amplitude (in milliamperes) endured for a single 45 mg food pellet, Punishment Effort Ratio was calculated as the ratio of footshock breakpoint to lever press breakpoint, and Delay Punishment Ratio was calculated as the ratio of the footshock breakpoint at a 40-second delay compared to a 0-second delay. From the runway operant cocaine seeking task, Run Start was measured as the delay between the door opening and the animal leaving the starting chamber, while Run Reversals were measured by counting the number of run direction reversals during the task. These continuous behavioral data were analyzed using linear models to identify and regress out effects of covariates explaining > 2% of variance in phenotype and quantile normalized. Additionally, the median latency to avoid cocaine in the runway operant cocaine seeking task was used to categorize animals as High Cocaine-Avoiding. A summary of phenotypes is included in Supplemental Table 1.

To test the strength of linear relationships between phenotypes, Pearson correlations were estimated. SNP-based heritability (*h*^2^) and genetic correlations (*r_g_*) between traits were estimated using Genome-Wide Complex Trait Analysis and Genomic-Relatedness-Matrix Restricted Maximum Likelihood (GCTA- GREML)^21,22^. Genetic correlations were then used as distance to calculate hierarchical clustering of phenotypes using the complete-linkage method.^23^

### Genome-Wide and Phenome-Wide Association Studies

GWAS was performed using Mixed Linear Model Analysis with dosages and genetic-relatedness matrices using the leave-one-chromosome-out method implemented to account for relatedness in the HS Rat population. Significance thresholds were estimated by permutation analysis (*n*_permutations_ = 1000). This permutation analysis identified genome-wide significance thresholds of -log_10_(*p*) = 5.54 and 5.16 for *p* < 0.05 and *p* < 0.10, respectively. Quantitative trait loci (QTL) were identified by scanning each chromosome for associations with SNPs with *p*-values greater than permutation-derived thresholds. QTL intervals were determined by estimating linkage disequilibrium (LD) between SNPs using plink^24^ to identify SNPs with strong LD (*r*^2^ > 0.6) with the peak marker^25^. To identify additional loci on the same chromosome, the top SNP from the first QTL was used as a covariate and additional GWAS were performed until no additional SNPs with significant associations were identified on that chromosome.

To further prioritize candidate genes from QTL intervals, variant effects, eQTL, splice QTL (sQTL), and human GWAS associations were identified. To determine variant effects, variant annotations were performed using SnpEff^26^. To identify eQTL and sQTL for genes within GWAS QTL intervals, data from RatGTEx (RatGTEx.org) were used^27^. To identify associations with related traits from human GWAS, human orthologs of genes within QTL intervals were queried on GWAS Catalog (ebi.ac.uk/gwas)^28^.

A phenome-Wide Association Study (PheWAS) was employed to identify relationships between the identified genetic loci and other phenotypes identified in other HS rat cohorts^29^ from the Center for GWAS in Outbred Rats Database (C-GORD, RRID: SCR_021866; now available through ratgenes.org). PheWAS was conducted for variants with a linkage disequilibrium score *r*^2^ < 0.6 within a 3-Mbp window of the peak SNP from 14 suggestive (*p* < 0.1) or significant (*p* < 0.05) GWAS loci.

## Results

### Heritability and Correlations

Heritability estimates for behavioral phenotypes ranged from 0.0 (for both Run Reversals in females and binary High Cocaine Avoid in males) to 0.307 (for Run Reversals across both sexes), with Body Mass at 0.347 (Figure 1B, Supplemental Table 2). The binary High Cocaine Avoid traits were not significantly heritable in either sex or in both sexes combined, Run Reversals were not significantly heritable in females, and all other traits were significantly heritable.

**Figure 1.**
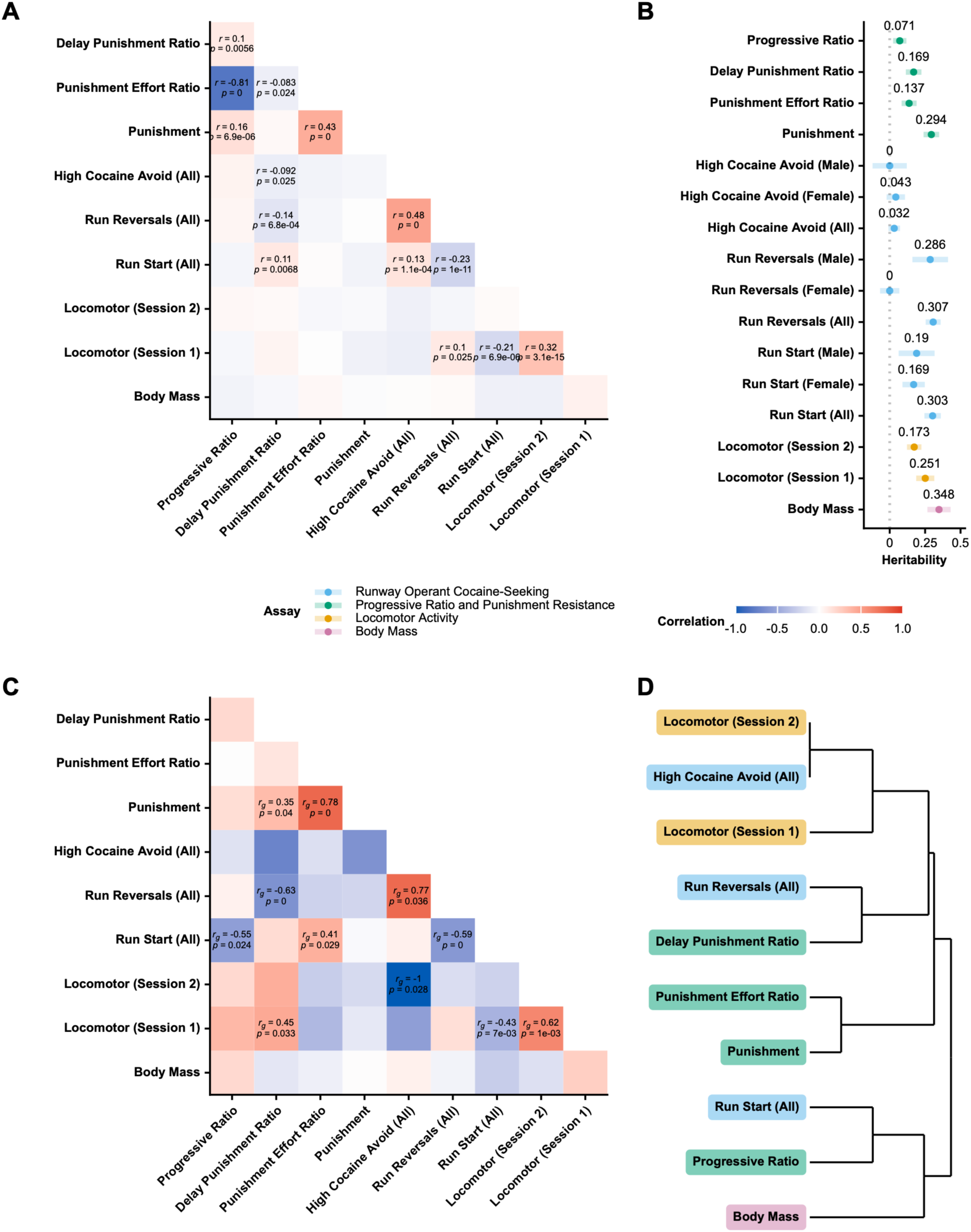
Heritability estimates and phenotypic and genetic correlations between cocaine avoidance and aversion behaviors. Pearson correlations were identified across phenotypes, identifying significant correlations across different behavioral assays including Delay Punishment Ratio and High Cocaine Avoid, Run Start, and Run Reversals (A). SNP-based heritability was estimated for phenotypic measures using GCTA-GREML (B). Significant heritability was estimated for all traits except Run Reversals in females and the binary High Cocaine Avoidance trait (in males and females separately as well as with both sexes combined). Exact heritability and correlation values are available in Supplemental Table 3. Genetic correlations were estimated using GCTA (C). Hierarchical clustering of traits was performed using genotypic correlations as distance using the complete- linkage method, identifying clusters of phenotypes from different behavioral assays such as Run Start and Progressive Ratio, Run Reversals and delay Punishment Ratio, or Locomotor (Session 2) and High Cocaine Avoid (D). These clusters suggest overlapping genetic architecture of aversion-based learning traits across different behavioral testing paradigms.

Significant phenotypic correlations were observed across different behavioral paradigms, such as Locomotor (Session 1) and Run Start (*r* = -0.206), Delay Punishment Ratio and Run Start (*r* = 0.111), Delay Punishment Ratio and Run Reversals (*r* = -0.139) (Figure 1A; Supplemental Table 3). Significant genetic correlations were also identified for phenotypes measured across different tasks (Figure 1C), including between Progressive Ratio from food-seeking task and Punishment (*r*_phenotype_ = 0.157; *r*_g_ = 0.175), Delay Punishment Ratio and Run Start (*r*_phenotype_ = 0.111; *r*_g_ = 0.178), and Locomotor (Session 1) and Run Reversals (*r*_phenotype_ = 0.104; *r*_g_ = 0.159), among others (Supplemental Table 3).

### QTL Identified for Punishment Task, Cocaine Avoidance, and Locomotion

A total of eight significant (*p* < 0.05) and five suggestive (*p* < 0.10) quantitative trait loci (QTL) were identified for behavioral phenotypes (Figure 2A; Table 1), including four significant loci for Punishment on chromosomes 2 (Supplemental Figure 4B), 3 (Supplemental Figure 4C), 5 (Figure 2B), and 6 (Figure 2C). The Punishment QTL on chromosome 5 had the highest -log_10_(*p*) of any autosomal QTL at 7.189 (Figure 2B). A significant locomotor QTL was also identified on chromosome 5 [-log_10_(*p*) = 5.60] (Supplemental Figure 4A). Finally, three significant loci were identified on the X chromosome for High Cocaine Avoid in both sexes [- log_10_(*p*) = 7.48] (Supplemental Figure 2B), High Cocaine Avoid in females only [-log_10_(*p*) = 7.40] (Supplemental Figure 2C, and Run Start in males [-log_10_(*p*) = 6.19] (Supplemental Figure 2D).

**Figure 2.**
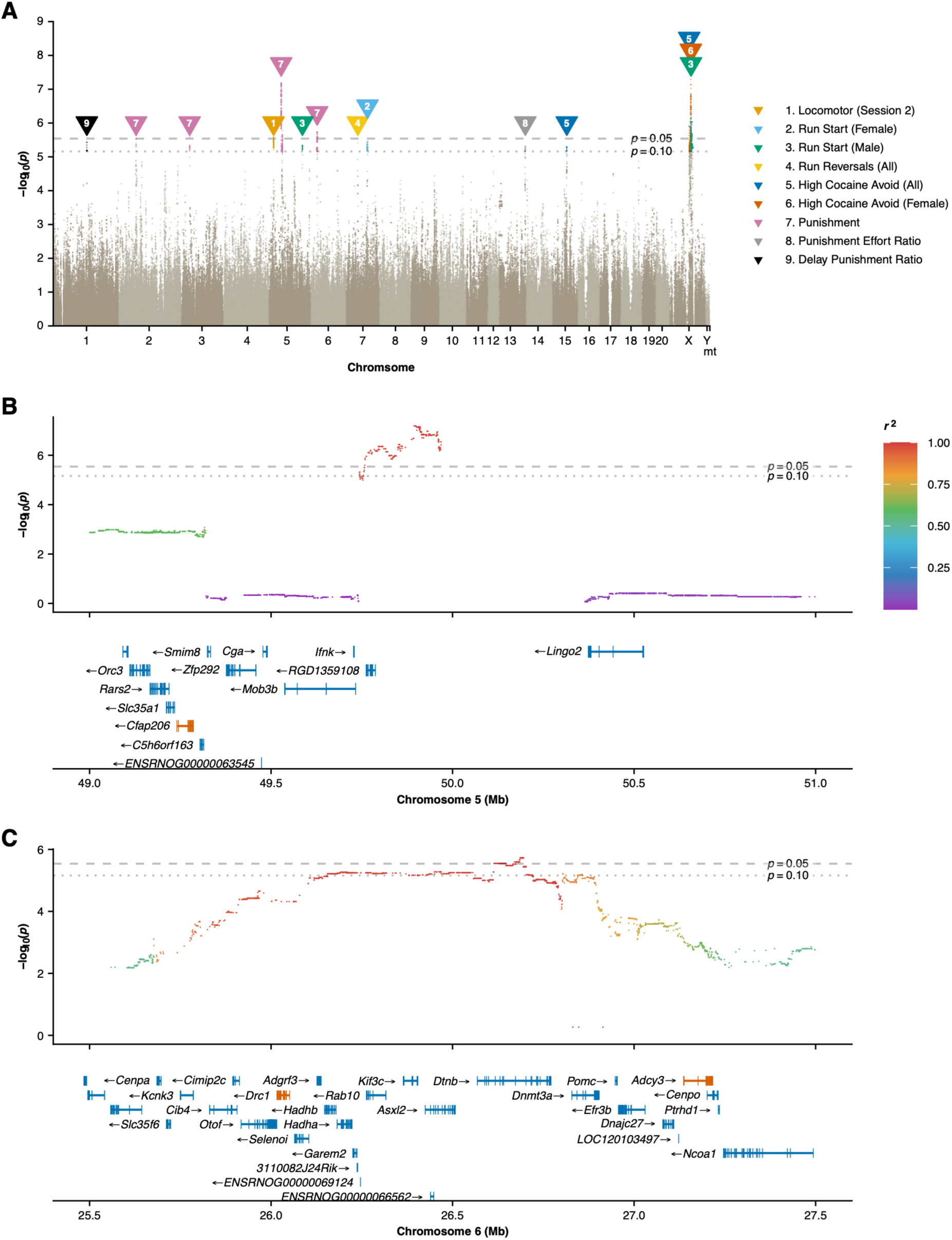
Loci related to aversion-based learning, cocaine avoidance, and locomotor activity identified in HS Rats. GWAS was conducted for individual traits, (A) with significant loci (*p* < 0.05) identified for Punishment (B and C), Locomotor activity in the first session, High Cocaine Avoidance, and Run Start in males only. GWAS was performed using Mixed Linear Model Analysis with dosages and genetic-relatedness matrices using the leave-one-chromosome-out method implemented to account for relatedness in the HS Rat population. Significance thresholds were estimated by permutation analysis (*n*_permutations_ = 1000). Suggested and significant loci are labeled numerically; heights of labels do not correspond with -log_10_(*p*) values. Significant loci for the Punishment trait were identified on chromosomes 5 (B) and 6 (C). Points are colored by linkage disequilibrium score *r*^2^ estimated by plink. Gene annotations were queried from Ensembl database mRatBN7.2. Highlighted genes include *Cfap206* (B), *Drc1*, and *Adcy3* (C), genes associated with primary neuronal cilia.

**Table 1.**
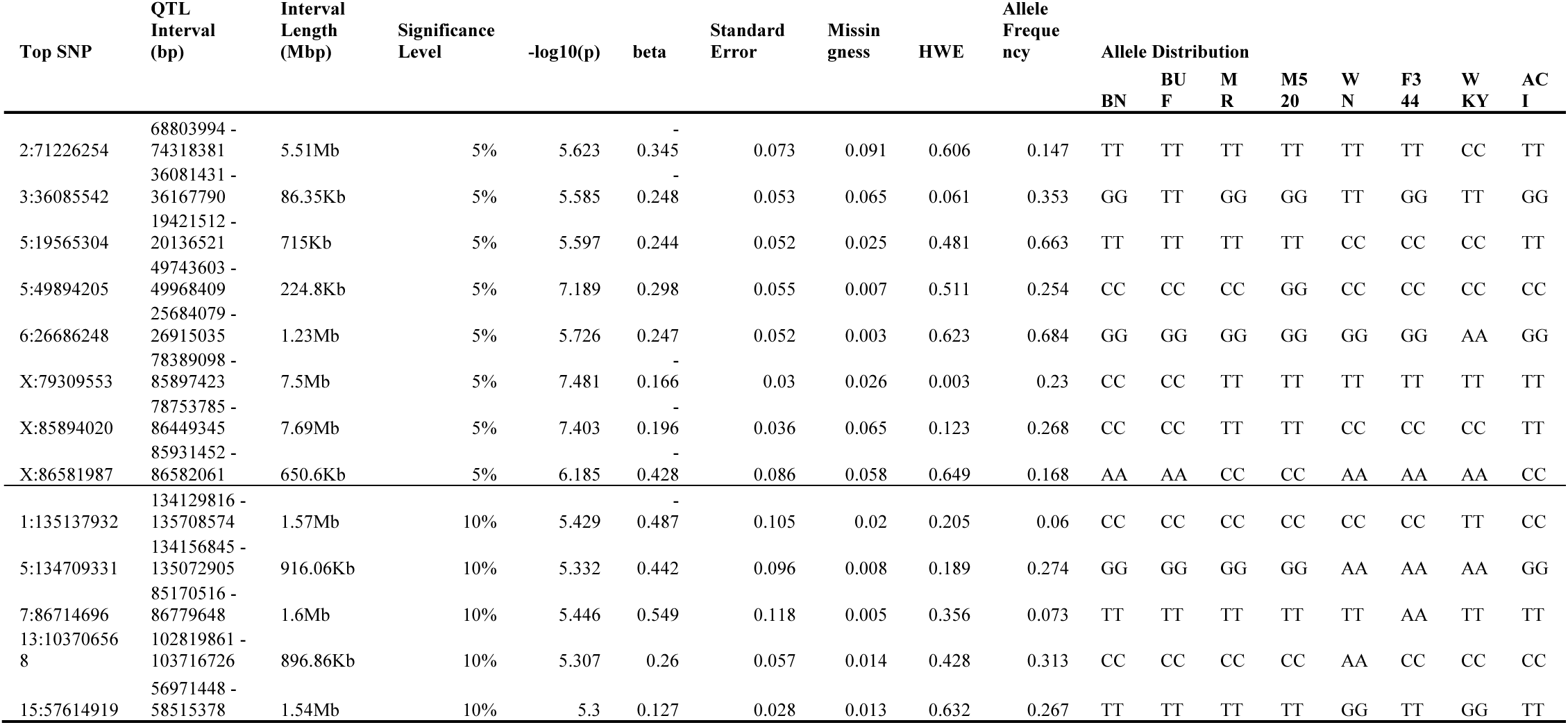
Significant (*p* < 0.05) and suggestive (*p* < 0.10) loci associated with avoidance-based learning traits in HS Rats. Quantitative trait loci (QTL) were identified by scanning each chromosome for associations with SNPs with *p*-values greater than permutation-derived thresholds (5.54 for *p* < 0.05; 5.16 for *p* < 0.10). QTL intervals were determined by estimating linkage disequilibrium (LD) between SNPs using plink to identify SNPs with strong LD (*r*^2^ > 0.6) with the peak marker.

### Candidate Gene Identification

A total of 172 genes, predicted genes, and pseudogenes were identified as positional candidates from significant or suggestive QTL, ranging from 0 to 38 genes per individual interval (Supplemental Table 4). The significant locus on chromosome 2 for Punishment contained a cluster of three cadherin family genes: *Cdh10*, *Cdh12*, and *Cdh18* (Supplemental Figure 3B). *Cdh10* has been associated with nicotine dependence^30^ and smoking initiation^31^ in human GWAS. *Cdh12* has also been associated with smoking initiation,^32–35^ as well as schizophrenia^36^ and neuroticism and cognitive function scores.^37^ *Cdh18* has been associated with alcohol consumption,^34,38,39^ smoking intiation,^39^ major depressive disorder,^40,41^ and verbal short-term memory.^42^ Of the three genes, only *Cdh12* has a *cis*-eQTL in the HS rat database, which was present in both whole brain and nucleus accumbens (Supplemental Table 6).

The Punishment QTL on chromosome 5 spans two genes and one predicted gene (Figure 2B). *Mob3b*, *Ifnk*, and RGD1359108 have not been implicated in addiction biology or other behavioral phenotypes in humans or animal models. Nearby genes outside of the QTL linkage disequilibrium interval include *Cfap206*, a gene involved in in brain development.^43^ *Cfap206* shows significant cis-eQTL in multiple brain regions (Supplemental Table 6).

The QTL on chromosome 6 for Punishment spans 1.23 Mbp and contains 29 genes (Figure 2C). Of these, four genes had coding variations in the HS Rat population: *Drc1*, *Dtnb*, *Asxl2*, and *Adcy3*, each having a missense variant. While not associated with any behavioral phenotype in human GWAS, *Asxl2* is associated with brain volume^44^ and white matter microstructure.^45^ *Adcy3* is associated with smoking initiation^39^ and cognitive function^46^, but also robustly associated with body mass index, body composition, and adiposity measures.^47–50^ Neither *Drc1* nor *Dtnb* had relevant associations in GWAS Catalog.

For the Locomotor (Session 2) locus on chromosome 5 (Supplemental Figure 4A), nine genes were identified. One gene in particular, *Car8*, has a known locomotor phenotype in mice^51^ and is important in cerebellar development, with variants known in humans to cause cerebellar ataxia.^51^ Another gene, *Nsmaf*, has a missense variant in the HS rat population (Supplemental Table 5), but is not typically associated with any locomotor phenotypes.

On the X chromosome, two overlapping loci for High Cocaine Avoid were detected (Supplemental Figure 2). In total, the overlapping loci contain 12 genes and 1 predicted gene. Of these, both *Klhl4* and *Dach2* were associated with memory decline in humans,^52^ but none of these genes are previously associated with addiction behaviors. The other X chromosome QTL for Run Start in males only also overlaps with the High Cocaine Avoid loci, with *Pcdh11x* the only gene shared by all three loci. *Pcdh11x* is a protocadherin gene in the same family as *Cdh10*, *Cdh12*, and *Cdh18* genes identified on chromosome 2 for the Punishment phenotype and regulates cell-cell adhesion in the nervous system as well as being implicated in development of cerebral asymmetry.^53^

### PheWAS

Variants associated with Punishment on chromosome 2 were also associated with reaction time in independent samples of HS rats (Supplemental Table 5).^54^ Other variants on the X chromosome were associated with novelty-seeking behavior^55^ and body mass in independent samples of HS rats. Variants from the chromosome 6 Punishment locus were also associated with body composition. Run Start in males and the High Cocaine Avoid phenotype loci on Chromosome X were also associated with locomotor phenotypes^56^ and social interaction phenotypes.^55^ No associations with other traits were observed for the Locomotor (Session 2) phenotype on chromosome 5.

## Discussion

We performed GWAS for several novel behavioral aversion-based learning paradigms which may contribute to the development of CUD. We identified eight significant associations (*p* < 0.05). Of these, cocaine-related phenotypes were only significantly associated with a single overlapping locus on Chromosome X whereas the food-based Punishment task was robustly associated with four significant loci; thus, 50% of our significant findings were about food-based Punishment. While there are theoretical reasons to believe that food- and cocaine-based aversion-based learning paradigms are related, and we did identify limited genetic or phenotypic correlations between them, the individual loci we identified for punishment were not significant for any of the cocaine aversion behaviors.

Most avoidance learning-based phenotypes were significantly heritable (Figure 1B), except for the High Cocaine Avoid binary trait (*h*^2^ = 0.032; 95% C.I. = -0.048-0.112); however, the lack of variance in the binary trait may have led to decreased heritability estimates. Run Reversals were not significantly heritable in females but were heritable in both males alone (*h*^2^ = 0.286; 95% C.I. = 0.039-0.533) and in the sexes combined (*h*^2^ = 0.307; 95% C.I. = 0.203-0.411). The highest heritability estimates were for Body Mass (*h*^2^ = 0.384; 95% C.I. = 0.223-0.545) and Run Reversals in both sexes combined. The individual variation in these traits in our HS rat population and significant heritability estimates reinforce the value of study into avoidance-based learning as a component of addiction biology.

While genetic correlations across different behavioral paradigms largely did not reach the *p* < 0.05 level of significance (Supplemental Table 3), Run Reversals and Delay Punishment Ratio were inversely genetically correlated (*r_g_* = -0.632; *p* < 0.001), suggesting that similar genetic factors influence both cocaine avoidance and resistance to foot shock. Similarly, while High Cocaine Avoid and the two Locomotor sessions were not significantly genetically correlated, they clustered together in hierarchical clustering. Variants associated with the Punishment task were also associated with locomotor phenotypes as well as novelty-related behavior in other samples of HS Rats through PheWAS analysis. Novelty-seeking and locomotor behavior have both been routinely suggested as predictors of substance self-administration in rodent models^57–59^ and are hypothesized to be similar to risky, sensation-seeking, or externalizing behaviors related to SUD in humans.^60–62^

QTL on the X chromosome from both tasks (Figure 2A) overlapped, suggesting a shared genetic architecture for both the food-based punishment resistance assay and cocaine avoidance. GWAS identified several candidate genes -- particularly from autosomal QTLs for the Punishment task -- also implicated in substance use or other behavioral phenotypes in humans from the GWAS Catalog (Supplemental Table 3). Notably, all three cadhedrin genes *Cdh10*, *Cdh12*, and *Cdh18* associated with Punishment on chromosome 2 were also associated with initiation of smoking in humans, as was *Adcy3* from a separate Punishment locus on chromosome 5 (Figure 2A). These findings suggest a possible link between avoidance-based learning assayed in the Punishment task and addiction behaviors both including and extending beyond cocaine aversion.

Additional candidate genes identified from the significant locus for Punishment on chromosome 6 include *Cfap206* (Figure 2C) and *Drc1* (Figure 2B), which along with *Adcy3* suggest a role for neuronal cilia. *Adcy3* localizes in neuronal cilia, making it a useful marker for primary cilia in the brain.^63^ While these cilia do not possess synaptic structure, they have been implicated in cAMP signaling.^64^ *Cfap206* has been implicated in motile cilia formation in sperm and in brain development.^43^ *Drc1*’s human ortholog has been identified as the cause of ciliary dyskinesia.^65,66^ Cilia have long been studied for their roles in neurodevelopment^67,68^, but an emerging body of research suggests a link between primary neuronal cilia and regulation of behaviors related to substances including cocaine,^69^ alcohol,^70^ morphine,^71^ and amphetamine.^72^ However, most of these studies involve ablation of primary cilia in various neuronal cell types; further study would be necessary to identify mechanistically the relationship between neuronal cilia and these addiction-related traits.

Limitations of the study include an imbalanced sex ratio, which reduced power to detect significant loci in males and to detect sex-specific effects. Additionally, while both food and cocaine were used in different assays for avoidance-based learning, results may not be fully generalizable to other substances, particularly when the transition from initial rewarding effects to delayed aversive effects is more protracted. The use of food in the punishment avoidance assay also may also affect the chromosome 6 locus for Punishment, where one of the positional candidate genes (*Adcy3*) has been repeatedly and robustly associated with obesity and adiposity measures in both HS rats^73^ human GWAS.^74–79^ Neuronal cilia in the hypothalamus have been found to regulate adiposity and body weight, though these studies were not conducted using ablation of *Adcy3*, *Cfap206*, or *Drc1*.^80^ This locus may therefore be driven by genetic factors that influence the value of the food reward rather than the aversion to punishment. Nevertheless, dysregulation of reward circuits has been hypothesized to contribute to both food and drug motivation.^81^

Addiction biology is extremely genetically complex and incorporates multiple developmental stages and vulnerabilities in the progression from cocaine use to CUD. Most animal model studies focus on self-administration or dependence, but relatively few studies have attempted to model avoidance-based learning as it applies to the aversive effects of cocaine exposure. Our results suggest that these avoidance-based learning traits are themselves complex heritable traits which arise from multiple genetic loci and that these traits relate to other behavioral domains related to addiction biology, such as novelty-seeking and externalizing behavior.

## Acknowledgments

This work was funded by the National Institute on Drug Abuse (NIDA) grant 5U01DA044468-04 and National Institute on Alcohol Abuse and Alcoholism (NIAAA) T32 AA013525.

---

**Supplemental Figure 1.**
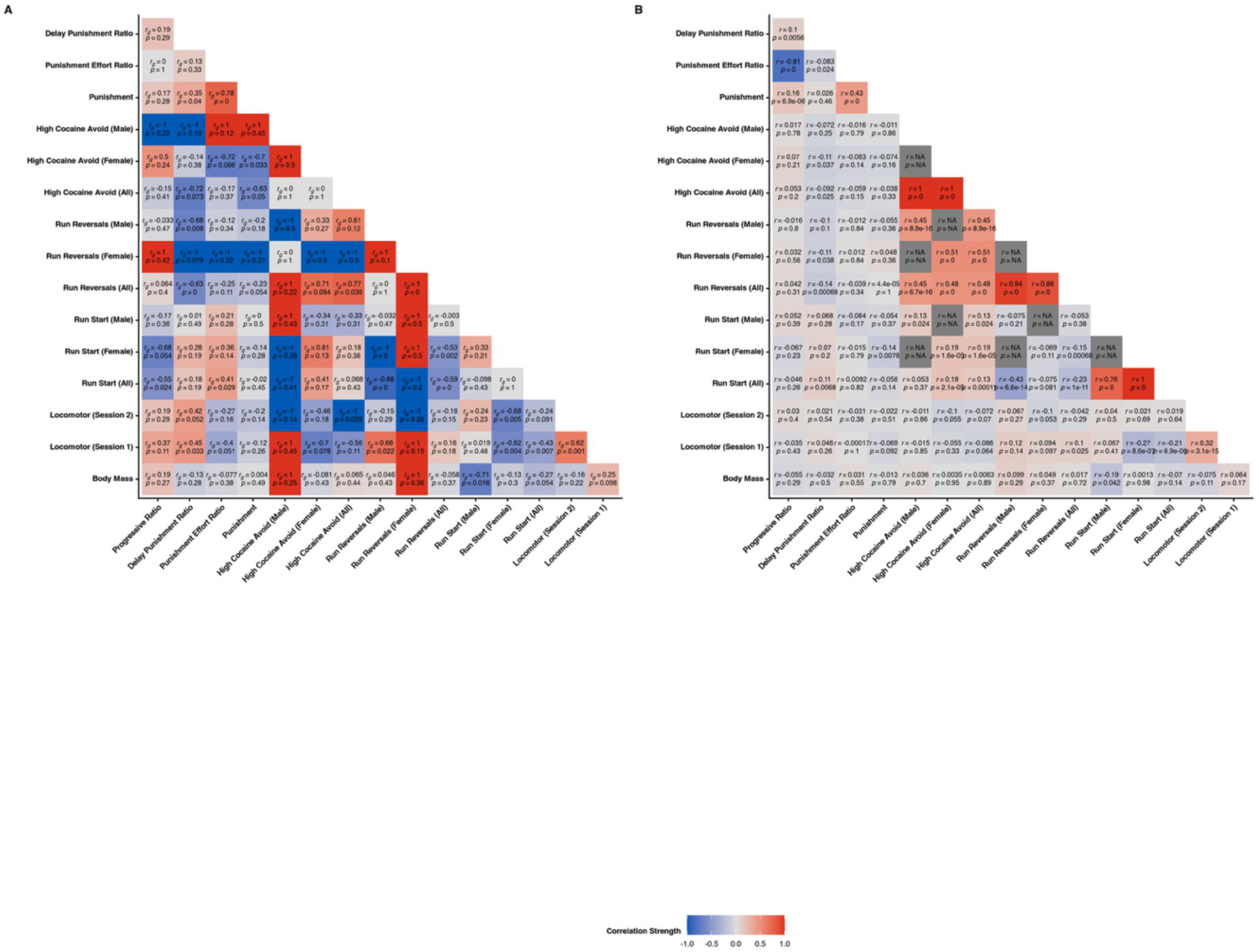
Phenotypic and genetic correlations across traits. Genetic correlations were estimated using GCTA (A), while Pearson correlations were used for phenotypes (B).

**Supplemental Figure 2.**
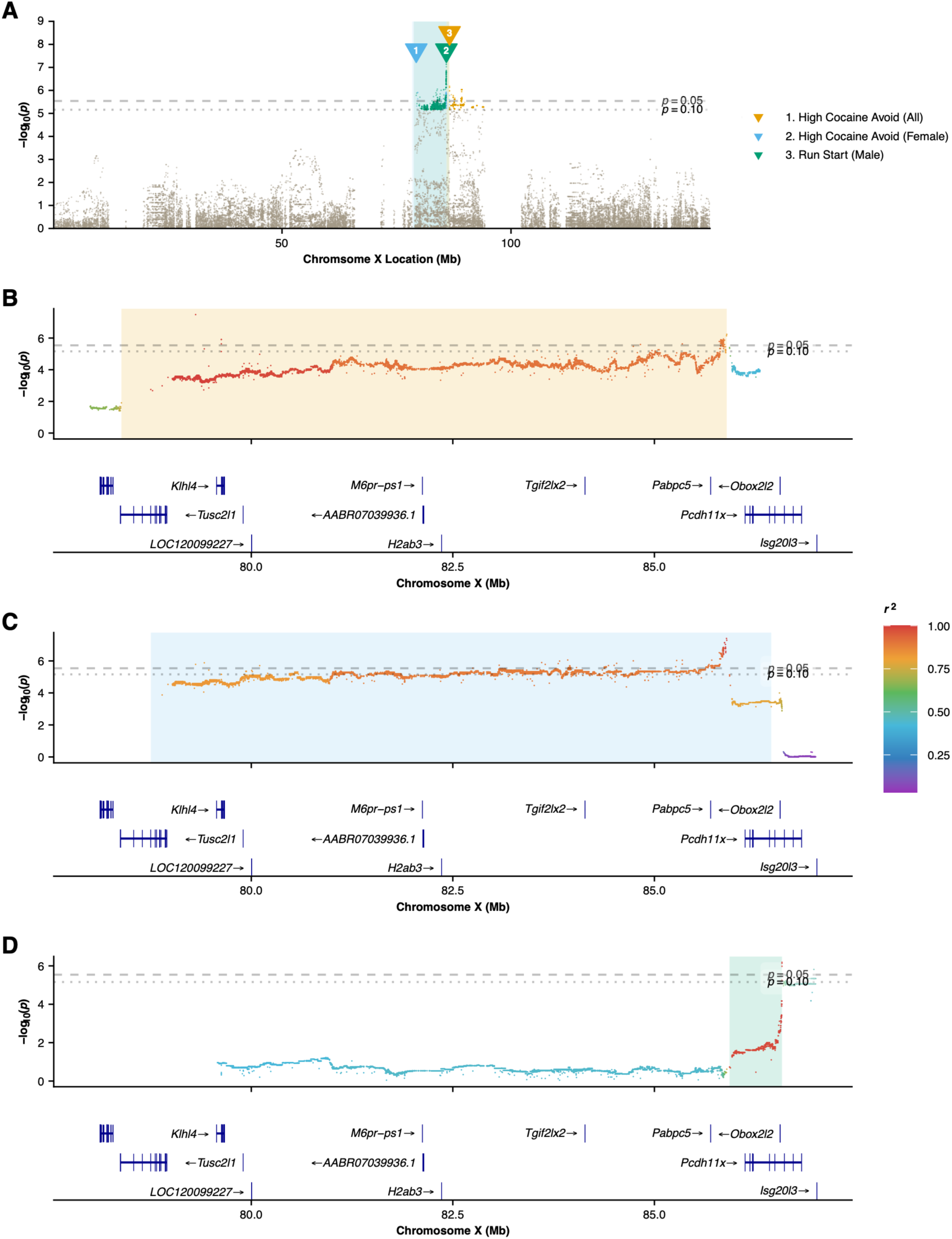
Loci identified on the X chromosome for High Cocaine Avoid (All), High Cocaine Avoid (Females), and Run Start (Males) overlap. Three significant (*p* < 0.05) quantitative trait loci were identified on the X chromosome from the runway operant cocaine-seeking task (A). Shaded regions indicate linkage disequilibrium intervals for identified QTL (*r*^2^ > 0.6). Loci are labeled numerically; heights of labels do not correspond with -log_10_(*p*) values. One locus was identified for High Cocaine Avoid (All) [X:79309553, - log_10_(*p*) = 7.581] (B), one for High Cocaine Avoid (Females) [X:85894020, -log_10_(*p*) = 7.403] (C), and one for Run Start (Males) [X:86581987, 6.185] (D). All three intervals overlap.

**Supplemental Figure 3.**
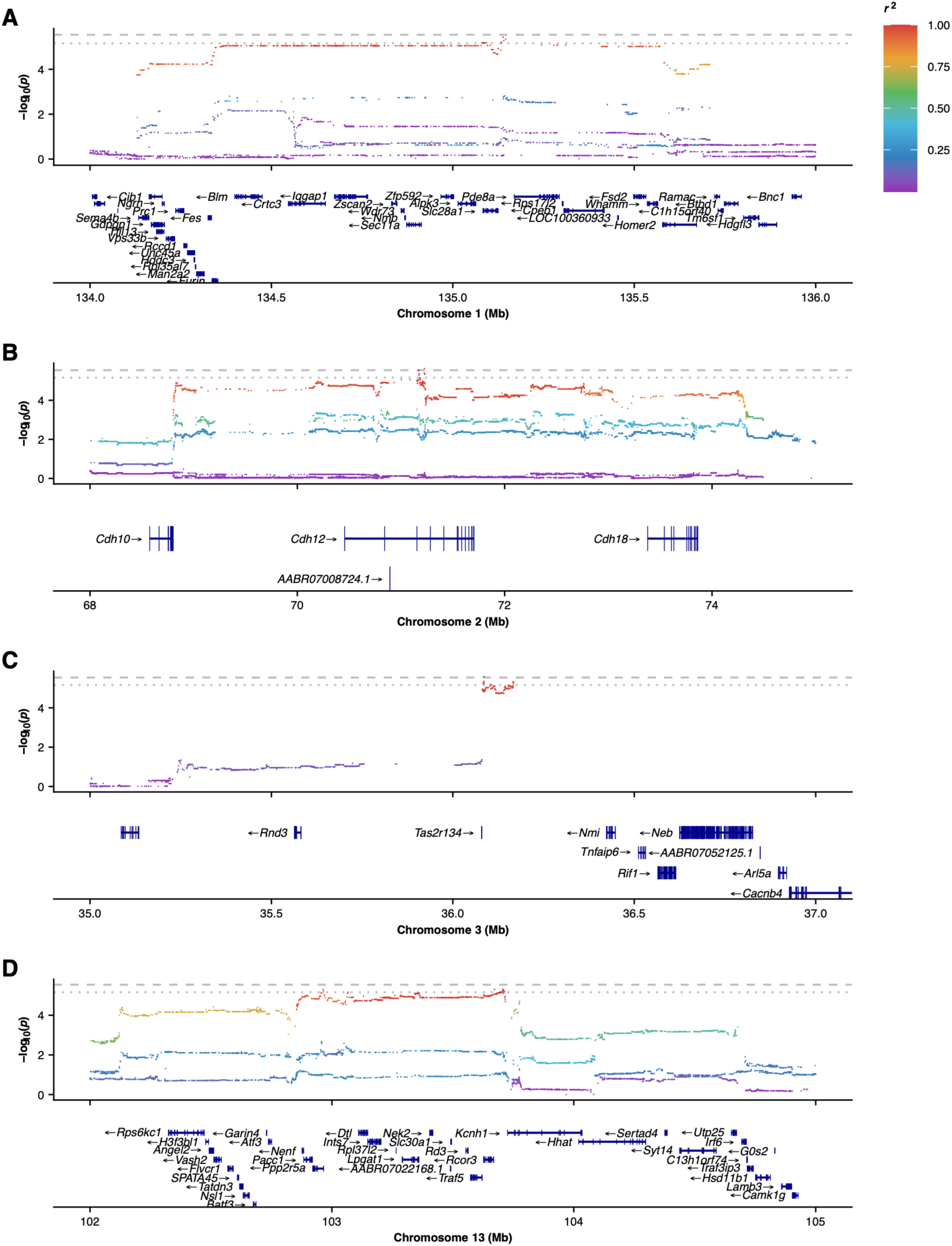
Additional significant and suggestive loci from the Progressive Ratio and Punishment Resistance tasks. A suggestive (*p* < 0.1) quantitative trait locus was identified for Delay Punishment Ratio on Chromosome 1 [1:135137932, -log_10_(*p*) = 5.429] (A). Two additional significant (*p* < 0.05) loci were also identified for Punishment, one on Chromosome 2 [2:71226254, -log_10_(*p*) = 5.429] (B) and one on Chromosome 3 [3:36085542, -log_10_(*p*) = 5.429] (C). One suggestive locus was also identified on Chromosome 13 for Punishment Effort Ratio [13:103706568, -log_10_(*p*) = 5.429] (D). Dashed lines represent a significance cutoff of *p* < 0.05; dotted lines represent *p* < 0.1.

**Supplemental Figure 4.**
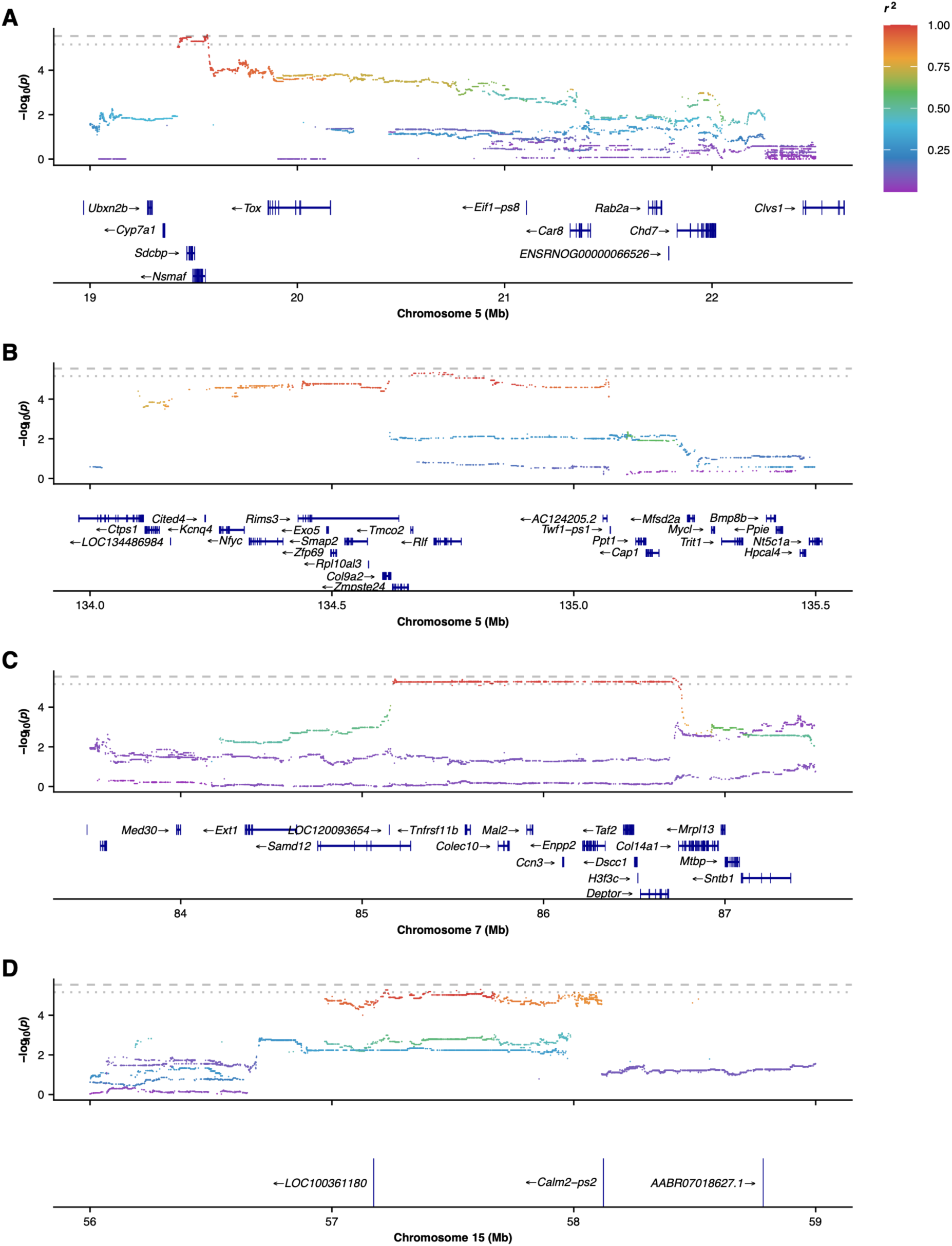
Additional significant and suggestive loci for Locomotor and Runway Operant Cocaine-Seeking tasks. A significant (*p* < 0.05) quantitative trait locus was identified for Locomotor (Session 2) on Chromosome 2 [2:71226254, -log_10_(*p*) = 5.623] (A). Suggestive (*p* < 0.1) loci were identified for Run Start (Males) on Chromosome 5 [5:134709331, -log_10_(*p*) = 5.332] (B) and Run Start (Females) on Chromosome 7 [7:86714696, -log_10_(*p*) = 5.446] (C). A suggestive locus was also identified for High Cocaine Avoid (All) on Chromosome 15 [15:57614919, -log_10_(*p*) = 5.300] (D). Dashed lines represent a significance cutoff of *p* < 0.05; dotted lines represent *p* < 0.1.

**Supplemental Table 1.** Phenotypes with brief descriptions and behavioral assays.

**Supplemental Table 2. Heritability estimates for phenotypes.** Narrow-sense (i.e., SNP-based) heritability was estimated using GCTA-GREML and animals’ genetic relatedness matrix.

**Supplemental Table 3. Genetic and phenotypic correlations.** Phenotypic correlations were generated using Pearson correlations in R. Genetic correlations were calculated using GCTA.

**Supplemental Table 4. Positional candidate genes from QTL intervals.** QTL intervals were determined by estimating linkage disequilibrium (LD) between SNPs using plink to identify SNPs with strong LD (*r*^2^ > 0.6) with the peak marker. Positional candidate genes were identified using ENSEMBL gene annotations.

**Supplemental Table 5. Moderate- and high-impact coding variants within QTL intervals.** Positional candidate genes were identified using ENSEMBL gene annotations. Impact of coding sequence variants was determined using SnpEff.

**Supplemental Table 6. eQTL results from RatGTEx**. Expression QTL for genes within QTL intervals were identified from RatGTEx.

**Supplemental Table 7. sQTL results from RatGTEx.** Splice QTL for genes within QTL intervals were identified from RatGTEx.

**Supplemental Table 8. PheWAS results for peak SNPs from QTL intervals.** The association between the top SNP for a trait from this study and all other top SNPs from traits in the HS Rat Phenome Database that were mapped within a 3 Mb window and a *r^2^* above 0.6.

